# Comparative Analysis of Methods to Reduce Activation Signature Gene Expression in PBMCs

**DOI:** 10.1101/2023.07.31.549036

**Authors:** Lovatiana Andriamboavonjy, Adam MacDonald, Laura K. Hamilton, Marjorie Labrecque, Marie-Noёlle Boivin, Jason Karamchandani, Jo Anne Stratton, Martine Tetreault

**Author notes:** + These authors contributed equally: Lovatiana Andriamboavonjy, Adam MacDonald. Corresponding authors &. ^7^ Lead contact (Martine Tetreault).

## Abstract

Preserving the *in vivo* cell transcriptome is essential for accurate profiling, yet factors during cell isolation including time *ex vivo* and temperature induce artifactual gene expression, particularly in stress-responsive immune cells. In this study, we investigated two methods to mitigate *ex vivo* activation signature gene (ASG) expression in peripheral blood mononuclear cells (PBMCs): transcription and translation inhibitors (TTis) and cold temperatures during isolation. Comparative analysis of PBMCs isolated with TTis revealed reduced ASG expression. However, TTi treatment impaired responsiveness to LPS stimulation in subsequent *in vitro* experiments. In contrast, cold isolation methods also prevented ASG expression; up to a point where the addition of TTis during cold isolation offered minimal additional advantage. These findings highlight the importance of considering the advantages and drawbacks of different isolation methods to ensure accurate interpretation of PBMC transcriptomic profiles.

**Highlights:**

- Traditional room temperature isolation methods trigger activation signature gene expression in PBMCs, even when rapidly isolated, whereas 4°C isolation methods do not.
- Transcription and translation inhibitors and cold processing techniques reduce activation signature gene expression via shared mechanisms.
- PBMCs treated with transcription and translation inhibitors lose responsiveness to external stimuli.
- Cold isolation methods offer a suitable and inexpensive alternative to mitigate activation signature gene expression in PBMCs.

## Introduction

The rise of high-throughput sequencing technologies has transformed the use of transcriptomics and enabled the study of gene expression in unprecedented detail. However, concerns regarding the quality and accuracy of transcriptomic data have emerged. Analyses have uncovered the presence of *ex vivo* activation gene signatures resulting from cell isolation techniques across various tissues and organs^1–6^. These findings emphasize the importance of rigorous quality control assessments as well as offer insights into biases introduced during sample preparation. Indeed, once cells are removed from their *in vivo* environment, induced cellular stress responses can have wide-reaching impacts on RNA profiles, causing RNA landscapes to gradually lose *in vivo* relevance^2^.

Ongoing studies are addressing these concerns by attempting to implement standardized protocols, improving isolation techniques, and developing computational methods to account for potential artifacts^2, 7, 8^. In fact, there has been a shift to maintain cells and tissues in cold conditions during isolation to mitigate transcriptional changes^9^. However, not all tissues may be amenable to cold isolation. For example, peripheral blood mononucleated cells (PBMCs); the main immune cells of the body, are most commonly isolated at room temperature using density gradient centrifugation. The option of cold processing has been largely disregarded, possibly due to concerns about the effectiveness of cell separation and apprehensions regarding unforeseen consequences associated with cold processing methods, such as effects on survival. While these hypothetical limitations exist, it remains unclear what optimizations would be required to efficiently isolate PBMCs at cold temperatures, or whether cold processing offers any benefit to standard PBMC isolation protocols. Recently, Marsh et al. utilized another method to mitigate processing artifacts. Using transcription and translation inhibitors (TTis), they revealed that immune cells from the brain and blood are particularly vulnerable to *ex vivo* activation during enzymatic digestion at 37°C and that the inclusion of TTis such as anisomycin, triptolide, and actinomycin D reduces the expression of 9 activation signature genes (ASGs), including AP1 complex genes (*JUNB*, *JUN*, *FOS*), as well as *DUSP1* (dual specificity phosphatase-1), *NFKB1A* (NF-kappa-b alpha subunit), *ZFP36* (zinc finger protein-36), *DDIT4* (DNA damage-inducible transcript-4), *CXCR4* (CXC-motif chemokine receptor-4), and *CCL4* (chemokine ligand-4)^10^.

Here, we examine the effects of these ASG-mitigating methods; TTis and cold processing on the transcriptome of PBMCs. We show that TTis indeed do reduce ASG expression caused by standard RT processing, however they also block the ability of PBMCs to respond to stimuli thereafter. On the other hand, we reveal that using a cold isolation protocol similarly dampens ASG expression and that the addition of TTis during cold isolation offers minimal additional advantage. Thus, while both TTis and cold isolation effectively reduced ASG expression, TTis should be used with caution if PBMCs are anticipated to be used for future functional assays.

## Results

### Transcription and translation inhibitors reduce activation signature gene expression

We first determined if ASG expression is regulated in PBMCs when using a traditional isolation protocol compared to that of PBMCs collected with a TTi cocktail (all at RT with RBC lysis) as described by Marsh et al. (Fig 1A). We performed bulk RNA sequencing on PBMCs, and found a total of 293 DEGs, of which the vast majority were down-regulated (Fig 1B). We also compared multiple conditions with and without RBC lysis buffer to determine if this influenced the DEGs. Interestingly, regardless of how PBMCs were isolated, either at RT (Fig. S1 A-C) or in cold conditions (Fig. S1D-F), with or without RBC lysis buffer, we observed few significant differentially expressed genes (DEGs); 28 DEGs at RT (Fig. S1B), 45 DEGs in cold conditions (Fig. S1E). Intriguingly, when we made the same comparisons, but under TTis conditions, we observed a large number of DEGs (274 DEGs) but only in the RT condition (Fig. S1C) versus 21 DEGs with cold (Fig. S1F). This suggests that the addition of TTis at RT triggers a transcriptomic response that is inhibited when PBMCs are processed cold or when RBC lysis buffer is added. Next, enrichment analysis was performed on the 280 DEGs decreased by TTis. Kyoto encyclopedia of genes and genomes (KEGG) revealed enrichment in pathways related to cell contact, apoptosis, and inflammatory IL-17 signaling following addition of TTis (Fig 1C). Gene ontology (GO) analysis for cellular compartment (CC) found strong enrichment of AP-1 complex genes including *FOS*, *JUNB*, and *JUN* genes, likewise identified by Marsh as well as cytoskeleton-related pathways (Fig 1D). Also in line with the Marsh findings, we observed reduced expression of their proposed ASG set following addition of TTis, with all but *DDIT4* being significantly down-regulated (Fig 1E).

**Figure 1:**
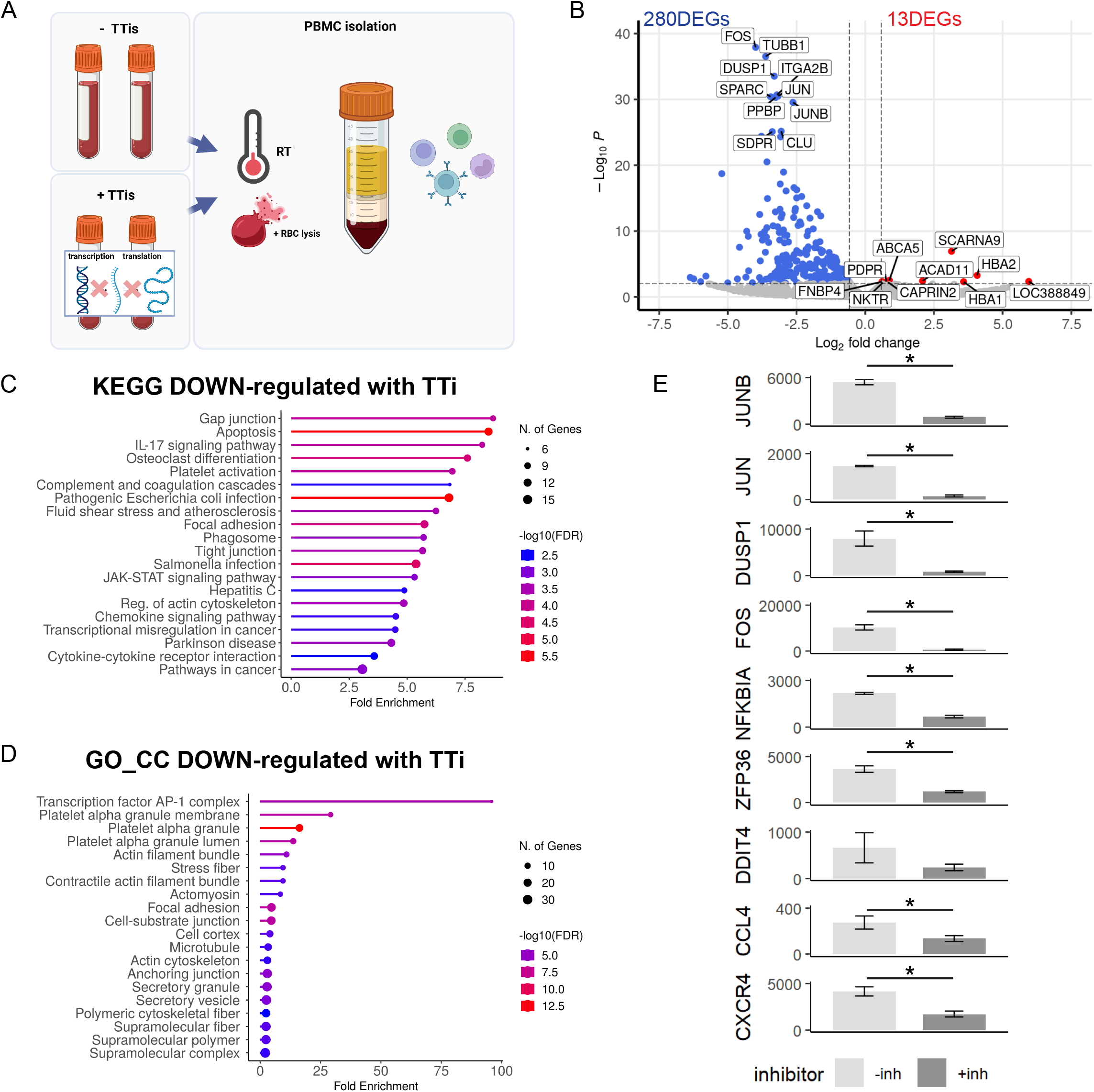
Transcription and translation inhibitors reduce activation signature genes. **A** Schematic of the experimental plan comparing the transcriptomic signatures from PBMCs isolated using a traditional protocol and PBMCs treated with and without a TTi cocktail (n=2/condition). **B** Volcano plot of 293 DEGs identified in the TTi-treated PBMCs compared to the traditional protocol. **C** Enrichment analysis of the 280 down-regulated DEGs revealed enrichment in KEGG pathways related to cell contact, apoptosis, and inflammatory IL-17 signaling. **D** Gene ontology (GO) enrichment analysis for cellular compartment (CC) showed strong clustering of AP-1 complex genes (*FOS*, *JUNB*, and *JUN*) and genes involved in multiple cytoskeleton-related pathways. **E** Bar graphs of the activation signature gene (ASG) set identified by Marsh et al., Error bars represent mean +/- SEM. Significance was set at p-value < 0.01 and an absolute log2FoldChange value of 0.58 (1.5 fold). Refer to TableS1 for raw transcriptomic data and TableS2 for complete DEG lists.

This data demonstrates that ASG expression in PBMCs is triggered by a traditional isolation protocol at RT, even with rapid isolation, and addition of TTis significantly reduces this activation.

### PBMCs treated with TTi during the isolation protocol no longer respond to stimuli *in vitro*

Considering that TTis essentially freeze gene expression, we next asked whether cells isolated in the presence of TTis would remain responsive to subsequent stimulation *in vitro*. PBMCs isolated with and without TTis were treated with the pro-inflammatory stimulus LPS for 6 hours, either directly after isolation (Fig 2A-C) or 18 hours after (Fig 2A, D, E). The gene expression of immune effectors *IL1β* and *IL6* indicated that PBMCs not exposed to TTi responded sharply to LPS, exhibiting increased expression of *IL1β* (Fig 2B, 2-way ANOVA Sidak’s post hoc, p=0.0256) and *IL6* (Fig Cc, 2-way ANOVA Sidak’s post hoc, p=0.0816) when exposed to LPS immediately after plating, as well as 18 hours after; IL1β (Fig 2D, 2-way ANOVA Sidak’s post hoc, p=0.0043) and IL6 (Fig 2E, 2-way ANOVA Sidak’s post hoc, p=0.0011). However, PBMCs treated with TTi during isolation showed no increase in *IL1β* (Fig 2B, 2-way ANOVA Sidak’s post hoc, p>0.9999) or *IL6* (Fig 2A, C, 2-way ANOVA Sidak’s post hoc, p>0.9999) when treated either immediately after plating or 18 hours later (IL1β: Fig 2D, 2-way ANOVA Sidak’s post hoc, p=0.9858; IL6: Fig 2E, 2-way ANOVA Sidak’s post hoc, p=0.9761), indicating complete unresponsiveness to LPS.

**Figure 2:**
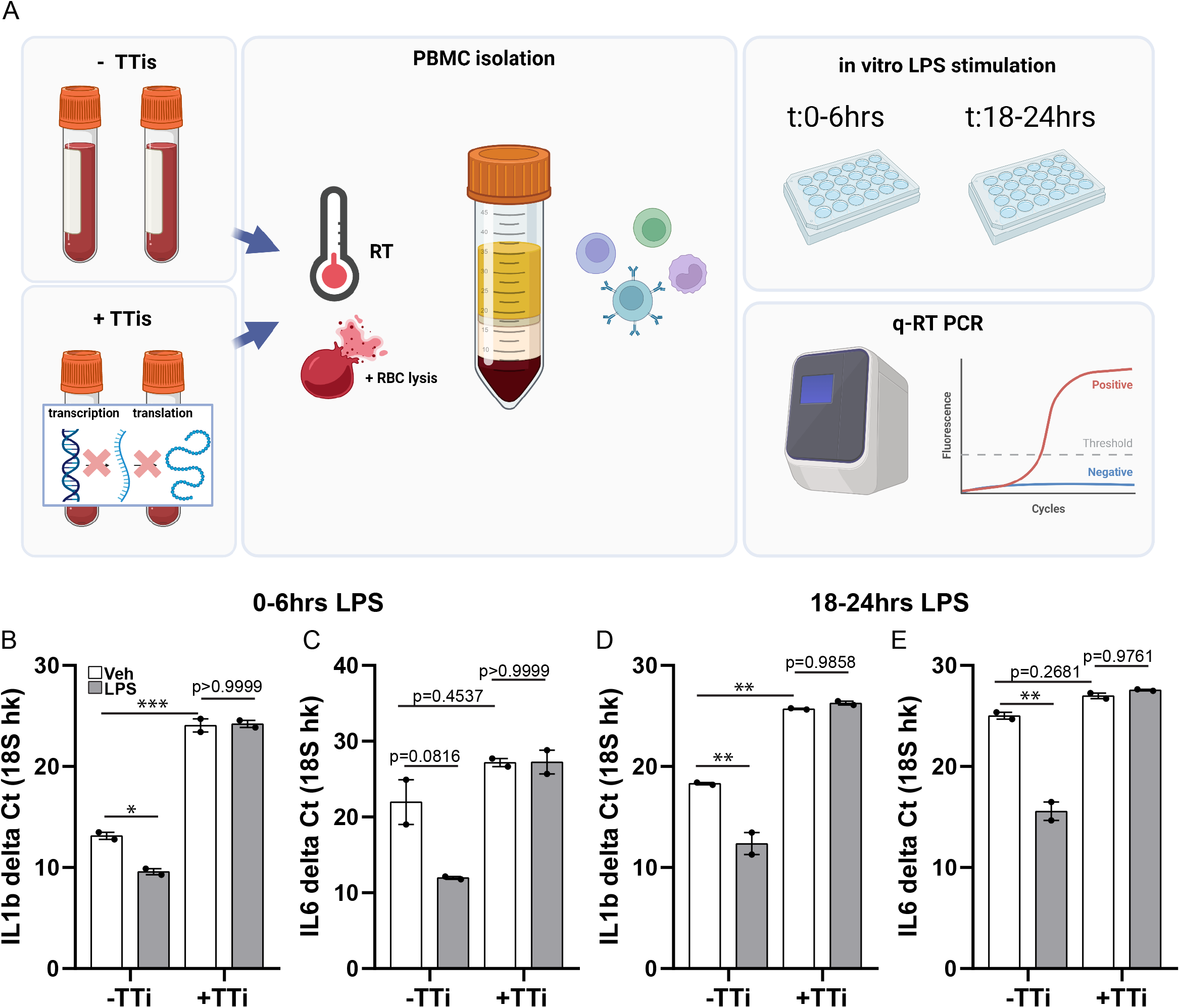
PBMCs treated with TTis during the isolation protocol no longer respond to stimuli *in vitro*. **A** Schematic of the experimental plan where PBMCs isolated with and without TTis were subjected to LPS stimulation for 6 hours either immediately after isolation or 18 hours post-isolation (n=2/condition). **B** Gene expression analysis of *IL1β* in PBMCs showed a significant increase in cells not exposed to TTi when treated with LPS at 0 hours post-treatment (p=0.0256, 2-way ANOVA Sidak’s post hoc), while no significant increase was observed in cells treated with TTi (p>0.9999, 2-way ANOVA Sidak’s post hoc). **C** Gene expression analysis of *IL6* in PBMCs exhibited a trend towards increased expression in cells not exposed to TTi (p=0.0816, 2-way ANOVA Sidak’s post hoc) when treated with LPS at 0 hours post-treatment, whereas no significant increase was observed in cells treated with TTi (p>0.9999, 2-way ANOVA Sidak’s post hoc). **D** Gene expression analysis of *IL1b* in PBMCs exhibited increased expression in cells not exposed to TTi (p=0.0043, 2-way ANOVA Sidak’s post hoc) when LPS stimulation was performed 18 hours after TTi treatment, no significant increase was observed in cells treated with TTi (p=0.9858, 2-way ANOVA Sidak’s post hoc). **E** Gene expression analysis of *IL6* in PBMCs exhibited increased expression in cells not exposed to TTi (p=0.0011, 2-way ANOVA Sidak’s post hoc) when LPS stimulation was performed 18 hours after TTi treatment, no significant increase was observed in cells treated with TTi (p=0.9761, 2-way ANOVA Sidak’s post hoc). Error bars represent mean +/- SEM. Significance was set at p-value < 0.05.

These results reveal that while TTis may offer advantages when RT dissociation is required, a repercussion is the loss of the cells’ ability to respond to stimuli *in vitro*.

### Cold isolation of PBMCs also down-regulates activation signature genes compared to RT

In other tissues, it is common practice to preserve *in vivo* gene expression states by keeping cells cold during isolation^11^. However, it has been assumed that maintaining cells at cold temperatures during PBMC isolation may not be feasible^12^. As the use of TTis is both costly and inflexible, our next objective was to isolate PBMCs while maintaining the workflow at 4°C (cold) compared to a traditional PBMC isolation protocol at RT (Fig 3A) to determine if we can use cold conditions to suppress ASG expression. RNA sequencing analysis revealed a total of 111 DEGs, with 38 up-regulated DEGs in cold samples and 73 down-regulated DEGs (Fig 3B). Pathway analysis of the down-regulated DEGs demonstrated enrichment in immune-related processes based on KEGG pathways (Fig 3C), while GO CC revealed a significant enrichment of AP-1 pathway genes, including *JUNB*, *JUN*, and *FOS* (Fig 3D). Intriguingly, maintaining cells in cold conditions significantly reduced the expression levels of all ASGs down-regulated by TTis, except for *CCL4* (Fig 3E). Collectively, this data demonstrates that PBMCs isolated cold exhibit low expression levels of ASGs compared to RT.

**Figure 3:**
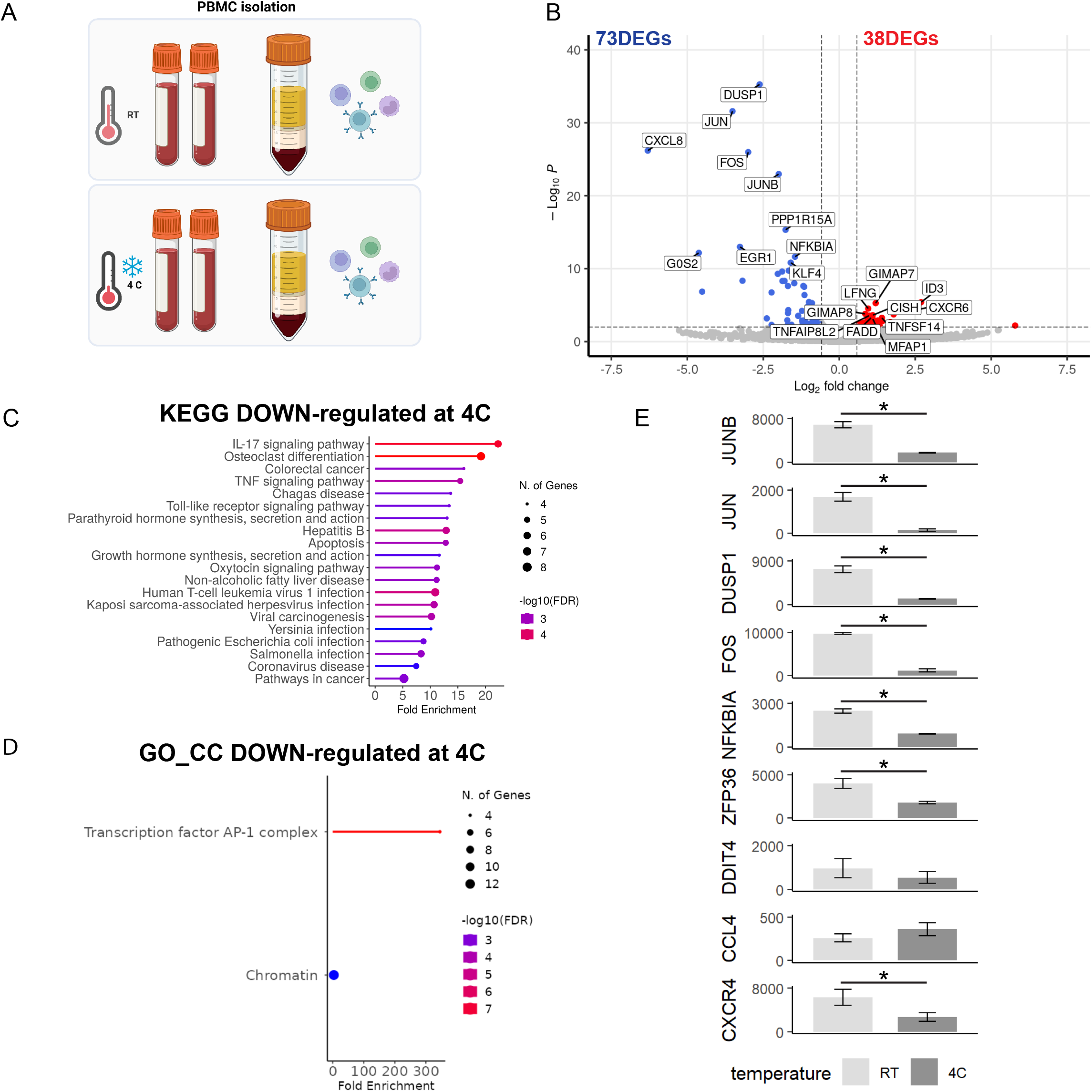
PBMCs processed cold compared to RT show down-regulation of activation signature genes. **A** Schematic of the experimental plan where PBMCs were isolated using a traditional protocol at either cold or room temperature (RT) (n=2/condition). **B** Volcano plot of RNA sequencing analysis revealing a total of 111 DEGs, with 38 up-regulated DEGs and 73 down-regulated DEGs in PBMCs isolated cold compared to RT. **C** KEGG pathway analysis of the down-regulated DEGs revealed enrichment in immune-related processes. **D** Gene ontology (GO) cellular compartment (CC) analysis demonstrated a significant enrichment of AP-1-stimulated pathway genes, including JUNB, JUN, and FOS (Fig 3d). **E** Bar graphs of the activation signature gene (ASG) set identified by Marsh et al., Error bars represent mean +/- SEM. Significance was set at p-value < 0.01 and an absolute log2FoldChange value of 0.58 (1.5 fold). Refer to TableS1 for raw transcriptomic data and TableS2 for complete DEG lists.

### Addition of TTi during cold PBMC isolation offers minimal transcriptomic advantage

Given the protective effect of keeping cells cold and the limitations imposed by the addition of TTi, the degree of transcriptomic similarity between PBMCs isolated cold and those processed with TTi was evaluated (Fig 4A). A comparison of the 157 DEGs decreased by cold processing (cold vs RT, all with RBC lysis buffer and without TTi) and the 280 DEGs decreased by the addition of TTi (+TTi vs -TTi, all with RBC lysis buffer processed at RT) revealed 64 DEGs shared between both processing methods (Fig 4B). Direct comparison of the expression of these 64 DEGs showed that most ASGs are similarly down-regulated by both methods; *JUN*, *DUSP1*, *FOS*, *ZFP36*, *NFKBIA*, *CXCR4*, and *JUNB* (Fig 4C). Pathway analysis of these shared DEGs predominantly identified immune-related KEGG pathways (Fig 4D), while GO CC revealed significant enrichment for AP-1 complex genes *JUN*, *JUNB*, *FOS*, along with platelet-, transcription-, and DNA-related compartments (Fig 4E). Intriguingly, 92 DEGs were selectively modified in cold samples, and a separate 216 DEGs were altered in TTi-processed samples. GO and KEGG pathway analysis of the DEGs specifically down-regulated by cold processing showed enrichment in ribosome-related processes (Fig S2A-B), while those down-regulated specifically by the addition of TTi were enriched in various processes, including platelets, cytoskeleton, and immune cell signaling (Fig S2C-D). To evaluate if keeping cells cold prevented *ex vivo* activation to the extent that the addition of TTi would no longer provide any benefit, we performed an additional experiment where TTi inhibitors were added, and cells were kept cold during the isolation protocol (Fig 4F). Interestingly, only 20 DEGs were observed, with 3 down-regulated and 17 up-regulated, when cells were harvested under these conditions. This indicated that the addition of TTi offers little advantage when cells are maintained in cold conditions during isolation (Fig 4G). Furthermore, the comparison of expression levels of the Marsh ASGs showed no significant differences between these conditions (Fig 4H).

**Figure 4:**
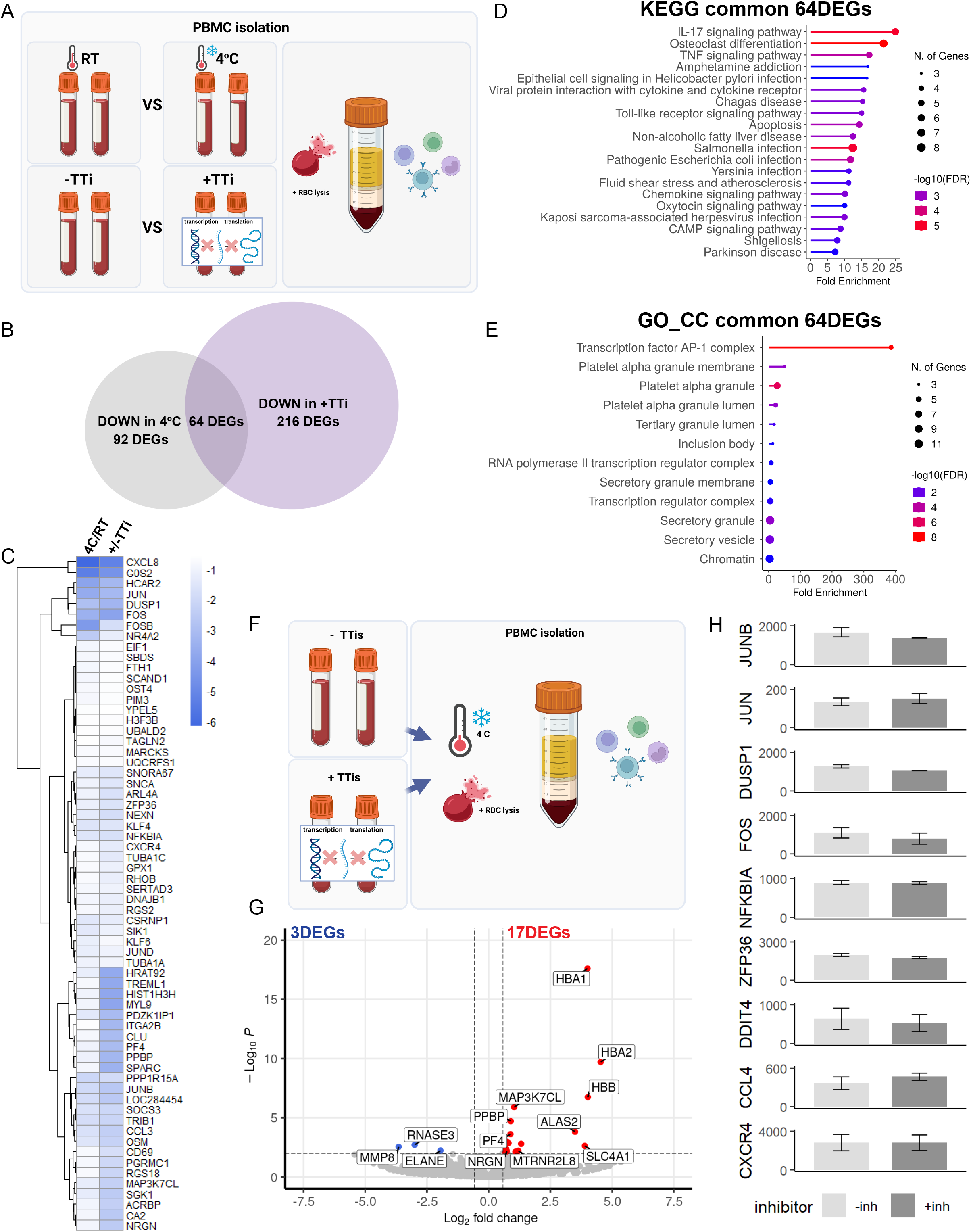
Transcription and translation inhibitors offer little transcriptomic advantage when a cold PBMC isolation protocol is used. **A** Schematic of the experimental steps to compare the degree of transcriptomic similarity between PBMCs isolated cold and those processed with TTi (n=2/condition). **B** Venn analysis of the differentially expressed genes (DEGs) decreased by cold processing and those decreased by the addition of TTi revealed 64 shared DEGs between both methods. **C** Heatmap of log2FC for shared DEGs decreased by cold processing or the addition of TTis. **D** KEGG pathway analysis of the shared DEGs predominantly identified immune-related pathways. **E** Gene ontology (GO) cellular compartment (CC) analysis revealed significant enrichment for AP-1 complex genes (*JUN*, *JUNB*, *FOS*) and other relevant compartments. **F** Schemati**c** of experiment conducted where TTi inhibitors were added, and cells were kept cold during the isolation protocol. **G** Volcano plot of DEGs when TTis are added to cold cell isolation showing 3 down-regulated and 17 up-regulated. **H** Bar graphs of the activation signature gene (ASG) set identified by Marsh et al., Error bars represent mean +/- SEM. Significance was set at p-value < 0.01 and an absolute log2FoldChange value of 0.58 (1.5 fold). Refer to TableS1 for raw transcriptomic data and TableS2 for complete DEG lists.

Collectively, this data illustrates a shared mechanism by both processing methods that converges to regulate multiple genes including AP-1 complex members.

### Thawed PBMCs show transiently increased activation gene expression *in vitro*

An advantage of PBMCs is their ability to be cryopreserved for subsequent experiments. However, how cryopreservation and time post thaw (pt) affect the transcriptome of PBMCs remains nebulous. Moreover, in an attempt to mitigate freeze/thaw-induced stress, some groups acclimate their cells *in vitro* after cryopreservation^13^, however we were unable to find any evidence on how time pt impacts ASG expression of PBMCs. In order to investigate this, we performed RNA sequencing on PBMCs immediately after thawing (t0pt) and compared them to PBMCs that were cultured for either 2 hours (t2pt) or 24 hours (t24pt) (Fig 5A-F). Global transcriptomic differences were observed among all three time points, as revealed by principal component analysis (Fig 5B). Comparing the transcriptome between t0pt and t2pt, 4978 DEGs were identified (2256 down-regulated, 2722 up-regulated). The number of DEGs between t0pt and t24pt totalled 2269 (1228 down-regulated, 1041 up-regulated) (Fig 5D), approximately half of what was observed between t0pt and t2pt. Interestingly, GO enrichment analysis of the down-regulated DEGs between t24pt and t0pt showed enrichment for AP-1 complex genes (*JUN*, *JUNB*, *JUND*, and *FOS*), along with vesicular trafficking-related compartments (Fig 5E). Analysis of ASG expression levels (Fig 5F) demonstrated that AP-1 complex genes (*JUNB*, *JUN*, *FOS*) and *DUSP1* decreased from t0pt to t2pt and continued to decline over 24 hours pt. In contrast, immune-related genes *(NFKB1A*, *ZFP36*, *DDIT4*, and *CXCR4*) either showed no change or significantly increased from t0pt to t2pt and sharply decreased by t24pt (Fig 5F). *CCL4*, however, exhibited an increase from t0pt to t2pt and remained elevated at t24pt (Fig 5F). To assess the kinetics of ASGs related to stress and immunity, we assessed how time pt affects stress signature (*JUNB*, *FOS*, *DUSP1*, *CCL4*) and cytokine genes (*IL1β*, *IL6*). We plated cryopreserved PBMCs in vitro for 0, 2, 6, 24, 48, and 72 hours (Fig 5G-H). We found that *JUNB* and *FOS* expression was similar at t0 and t2, then declined with time *in vitro*. *DUSP1* and *CXCR4* expression was approximately 4-to 8-fold higher at t2 compared with t0 and declined with time *in vitro*. *CCL4* was 9-fold higher at t2 compared to t0, increased until t6 then began to decline with time *in vitro* (Fig 5G). Expression of cytokines *IL1β* and *IL6* were approximately 16-fold higher at t2 from t0, peaked at t6, and declined thereafter (Fig 5H). This was consistent with the RNA sequencing data where *IL1β* and *IL6* were significant DEGs between t2pt and t0pt but not t24pt and t0pt (TableS2).

**Figure 5:**
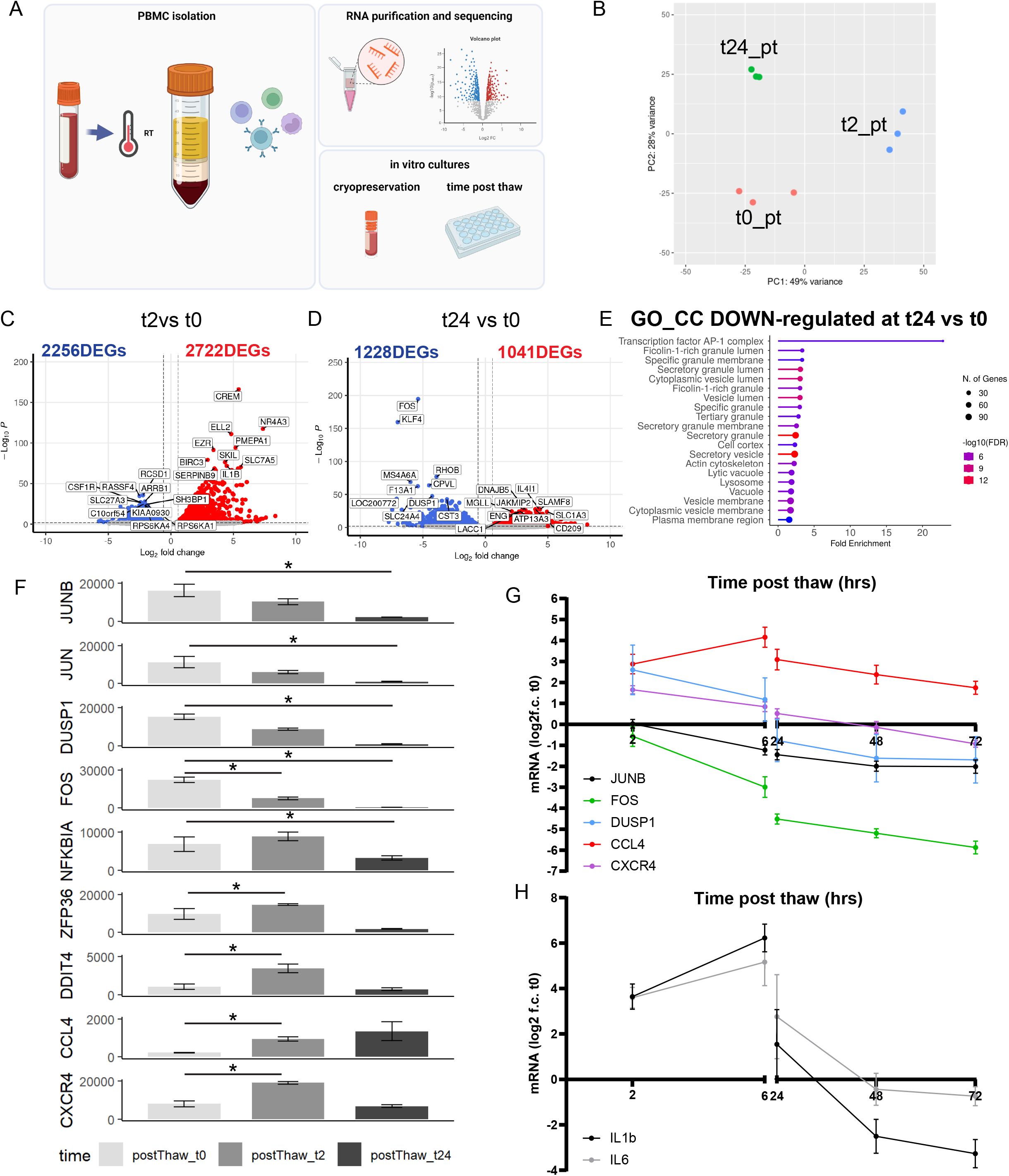
Cultured PBMCs show increased activation post thaw that dampens with time in vitro. **A** Schematic of the experimental plan for time post thaw analysis of cryopreserved PBMCs. **B** Principal component analysis revealed global transcriptomic differences among the three time points (n=2/condition). **C** Volcano plots of transcriptomic differences between 0 pt (t0pt) and 2 pt (t2pt) showing 4978 DEGs, with 2256 down-regulated and 2722 up-regulated. **D** Volcano plots of transcriptomic differences identified between t0pt and 24 pt (t24pt), with a total of 2269 DEGs (1228 down-regulated, 1041 up-regulated) **E** Gene ontology (GO) enrichment analysis of the down-regulated DEGs between t24pt and t0pt showed enrichment for AP-1 complex genes (*JUN*, *JUNB*, *JUND*, and *FOS*), along with vesicular trafficking-related compartments. **F** Bar graphs of the activation signature gene (ASG) set identified by Marsh et al., **G** Plot showing changes over time post thaw in mRNA expression of ASGs (*JUNB*, *FOS*, *DUSP1*, *CCL4*, and *CXCR4*) relative to their expression at t0pt (n=5/condition). **H** Plot showing changes over time post thaw in mRNA expression of immune genes (IL1b and IL6) relative to their expression at t0pt (n=5/condition). Error bars represent mean +/- SEM. Significance was set at p-value < 0.01 and an absolute log2FoldChange value of 0.58 (1.5 fold). Refer to TableS1 for raw transcriptomic data and TableS2 for complete DEG lists.

These findings highlight the impact of time post thaw on ASGs in cryopreserved PBMCs, with longer time *in vitro* (24-72 hours post thaw) having lowest expression of activation and immune-related genes.

## Discussion

In order to obtain comparable transcriptomic signatures, it is critical for the field to agree on a standardized method for processing PBMCs prior to analysis. Cold isolation protocols and TTis have both emerged as potential strategies for preserving *in vivo* cellular signatures^9, 10^. Cold isolation, by maintaining low temperatures during cell isolation, aims to minimize transcriptional changes and preserve the original gene expression profile. On the other hand, TTis halt transcription and translation processes, effectively freezing the cellular transcriptome and proteome at the time of treatment^10, 14^. In this article, we explore the positive and negative implications of each approach and highlight their shared mechanisms in order to inform decisions making for experimental designs.

### TTis effectively suppress ASG expression triggered during PBMC isolation

Recently, Marsh et al.^10^ demonstrated that TTis effectively suppress an *ex vivo* triggered response, which represents artificial changes in the transcriptomic profile of cells upon their removal from the body and that this suppression occurs through an AP-1-mediated mechanism. However, it should be noted that their study included additional mock digestion steps, including a 30-minute enzyme incubation at 37°C during PBMC isolation which are not necessary or commonplace in other labs. Thus, it remained unclear whether this signature was present in PBMCs isolated using traditional procedures. To investigate the practicality of TTis for PBMCs in a standard PBMC isolation protocol, we began by confirming that the TTis suppress ASG activation in PBMCs (Fig 1). In line with the findings of Marsh et al.^10^, we identified several down-regulated DEGs in samples treated with TTi vs those non-treated and very few upregulated DEGs. Among the most highly down-regulated genes are the signature genes identified by Marsh: *FOS*, *DUSP1*, *JUN*, and *JUNB*. Together, these genes drive “Transcription factor AP-1 complex” as the most significant GO term associated with the addition of TTis. This data demonstrates that TTis can effectively suppress genes that would otherwise become activated during standard PBMC isolation protocols.

### PBMCs no longer respond to stimuli *in vitro* following TTi treatment

Since TTis need to be added immediately once whole blood is collected, our next concern was that PBMCs treated with TTis may be irreversibly inhibited such that *in vitro* studies become infeasible. It would be ideal if these inhibitors could accommodate *in vitro* assays because if not, an entire vial of collected PBMCs would need to be dedicated solely to sequencing, thereby wasting cells that might otherwise be used to generate functional data^15^. To address this concern, we performed LPS stimulation either immediately or 18-hours after plating (Fig 2). This way, we could evaluate whether cells are immediately unresponsive following treatment with TTis, and if so, whether this effect is reversible. Surprisingly, in both conditions we found a complete lack of LPS response by PBMCs that were isolated with TTis. This reveals that isolation of PBMCs in the presence of TTis blocks their transcription and translation abilities for at least 18-24 hours, a key limitation.

### Cold processing is sufficient to suppress ASG expression in PBMCs

The widespread use of cold processing to minimize RNA degradation while slowing cellular metabolism for other cell and tissue types led us to next assess the feasibility of keeping PBMCs cold during isolation as an alternative to TTis (Fig 3). In comparing the DEGs between cold and RT processing, we observed the down-regulation of 73 genes in the cold condition, while 38 genes were upregulated (albeit with relatively lower significance compared to many down-regulated genes). Interestingly, genes down-regulated by this condition included the AP-1-related ASGs whose activation was similarly suppressed by addition of the TTis. Of the ASGs identified by Marsh et al.^10^, all showed either a statistically significant decrease or a trended decrease in gene expression; except for *CCL4* (Fig 3E). Kinetic evaluations of *CCL4* have previously revealed this gene to be expeditiously expressed following stimulation^16^, and as such it is likely that this gene activated prior to the cells attaining a cold temperature. Moreover, our time-course *post thaw* study showed that *CCL4* expression stays high for up to 72 hours *in vitro* (Fig 5G). Together, this data demonstrates that cold processing can be a viable strategy for suppressing ASGs in PBMCs.

### TTis and cold approaches act on shared regulatory mechanisms

To determine if TTis and cold processing share common regulatory mechanisms, we analyzed DEGs down-regulated by both methods. Intriguingly, we observed 64 shared genes; 41% of cold and 23% of TTi, which showed a 400-fold GO enrichment in AP-1 complex genes *FOS*, *DUSP1*, *JUN*, and *JUNB*. In addition, KEGG pathway analysis of the shared DEGs show enriched in inflammation-related processes including IL-17 and TNF signaling, as well as response to infection. Interestingly, comparing the levels of transcriptomic suppression of these common DEGs across methods showed that many are down-regulated to a similar degree (Fig 4C). We did observe two genes that are markedly more suppressed in the cold condition; C-X-C Motif Chemokine Ligand 8 (*CXCL8*) and G0/G1 Switch 2 (*G0S2*). *CXCL8* is involved in TNFα signaling^17^. The enhanced down-regulation of this gene in the cold condition likely represents a stronger anti-inflammatory effect being exerted on the PBMCs, yet this observation could also be an indicator of differential regulation of apoptosis^18^. Similarly, *G0S2* is known to be involved in enacting sudden metabolic shifts^19^, while also positively regulating apoptosis^20^, thus the down-regulation of this gene may carry anti-apoptotic effects as well. Of the DEGs that are commonly down-regulated, there is also a broad class that are more down-regulated in the TTi condition including; Heart tissue-associated transcript 92 (*HRAT92*), Triggering Receptor Expressed On Myeloid Cells Like 1 (*TREML1*), H3 Clustered Histone 10 (*HIST1H3H*), and Myosin Light Chain 9 (*MYL9*), which are implicated in a wide variety of cellular functions including transcriptional regulation, antigen presentation, histone assembly, and intracellular transport^21–24^. This hints at broad transcriptional down-regulation that is accentuated by TTis over standard or cold processing strategies, far beyond effectively reducing activation signature gene expression in PBMCs.

### Cold-induced changes in PBMC yield are unlikely to drive transcriptional changes

To preserve cell isolation integrity during density gradient centrifugation in our cold processed conditions we made the exception to keep the Ficoll at room temperature in all isolation conditions, however centrifugation steps and all other reagents were still at 4°C for the cold conditions. We qualitatively assessed the separation of the respective buffy coats while ensuring that consistent PBMC pellets were collected across the temperatures, noticing no significant differences. Moreover, trypan blue quantification revealed similar cell yield and viability. Within the bulk RNA sequencing data, we monitored GO terms to ensure that there weren’t any DEG clusters enriched for markers of specific cell types. In addition, many of the genes dysregulated by temperature were shared by the TTi condition, providing further confidence that these DEGs aren’t driven by temperature-dependent cell type differences. Altogether, we found no evidence that cold isolation altered cell proportions or yield to drive DEG expression.

### Time resting *in vitro* reduces ASGs after thawing but alters the transcriptome

Several groups culture the cells *in vitro* prior to downstream analysis^13^, we explored the transcriptomic effects of the temporal landscape experienced by PBMCs. We noted a significant change in approximately 5000 genes when comparing 2-hours *in vitro* against immediately harvested cells post thaw (t0pt). Similarly, there are over 2000 DEGs between 24-hours *in vitro* and t0pt. Intriguingly, we found that ASG expression levels generally decreased following 2-hours in culture, but that their expression levels fell to insignificant levels following 24-hours in culture. Overall, our data suggests that transcriptional changes tend to stabilize after approximately 24-hours *in vitro*, making this a more ideal time to compare expressional profiles across conditions. These results indicate that acclimating cells for 24 to 72 hours *in vitro* can reduce activation signatures. However, our findings also suggest that broad transcriptomic changes remain.

### Pros and cons of cold processing and TTis

When choosing between cold processing and adding TTis to PBMC processing protocols, consider that both methods effectively suppress gene expression quickly and efficiently. However, there are potential drawbacks to consider for each. Acute cold isolation protocols are effective at diminishing ASG expression, however, maintaining blood at cold temperatures prior to PBMC isolation for greater than 6-hours has been shown to reduce PBMC viability and influence purity^7^. TTis can also have unintended consequences on cellular functions beyond gene expression. For example, we observed 216 DEGs selectively down-regulated by TTis (Fig S2C-D) which were enriched for pathways related to red blood cells and cytoskeleton dynamics and that were not altered by cold processing. By blocking protein synthesis, TTis can disrupt normal cellular processes, potentially impacting cell viability, proliferation, and metabolism. Moreover, as we show here, TTis can interfere with the ability of cells to respond to stimuli, impairing their capacity to mount appropriate immune or stress responses. This limitation is particularly relevant when studying PBMCs, as their responsiveness to external signals is often critical for downstream experiments.

### Recommendations for rapid PBMC processing

It is important to recognize that our study processes PBMCs immediately. In cases where the whole blood is collected and then left idle for a period of time prior to PBMC isolation, we do not recommend leaving the whole blood cold^7^ or for long periods of time^25^, as previous work has shown that this can result in changes in gene expression, reduced viability and contamination with non-PBMC cell types. We strongly urge prioritizing the immediate PBMC processing in order to minimize *ex vivo* changes. In situations where it is desirable to block transcriptional changes immediately in whole blood, where cells can not be processed immediately, and where there is not an interest in observing the cells’ *in vitro* performance, it may be sensible to treat the whole blood with TTis. Importantly, we observed a previously unreported hemolytic effect caused by the TTi administration and internal testing showed that it was necessary to predilute the TTi in at least 500μl of PBS per 10ml of whole blood in order to quell the hemolysis. In other contexts, isolating the PBMCs cold can be a viable alternative: easier, less expensive, and more flexible for downstream applications.

In summary, while RNA sequencing has provided powerful insights into gene expression, it has also revealed shortcomings in our cell isolation methods. Finding optimal protocols to best preserve the *in vivo* cellular state is crucial for obtaining high-quality and biologically meaningful results. By comparing the ability of two isolation methods to mitigate *ex vivo* gene activation, we found that while TTis are effective for suppressing *ex vivo* ASG expression, their cost and inflexibility generally supports the use of alternative strategies for suppressing ASG expression in PBMCs, such as by processing the cells cold. Importantly, adding TTis to a cold isolation offered limited advantage. Together, our findings highlight the importance of selecting appropriate cell isolation methods to favour accurate biological profiling in studies involving immune cells.

## Materials and Methods

### Ethics statement

All experiments were approved by the Research Ethics office of the CRCHUM (2021-91150) or McGill University Research Ethics Office (IRB) of the Faculty of Medicine and Health Sciences (2021-6588).

### PBMC isolation

For Fig 1-4, experiments were performed on n=2 from blood from subjects; M;64 years and M;47 years. For Fig 5, sequencing experiments were performed on n=3 from blood from subjects M;78years, M;69years, F;66years and an additional n=2 F;69years, F;63years were analyzed for (Fig5 G-H).

For Fig 1-4 experiments, Human PBMCs were obtained by drawing blood into lithium heparin-coated tubes (cat#3048749) and for Fig 5 in BD vacutainer EDTA tubes (cat#366643). The blood collection followed the approved protocol of CRCHUM or McGill University Institutional Review Board, and informed consent was obtained from the participants. PBMC isolation began less than 10 minutes after the blood was drawn from the body. Briefly, peripheral blood samples were diluted with an equal volume of PBS containing 2% EDTA. Ficoll-Paque (cytiva, cat#17144003) was then underlaid, and centrifugation was performed at 800 × g for 20 minutes at room temperature (unless stated otherwise) to separate the middle white phase (buffy coat), which was transferred to a fresh tube. In cases where RBC lysis was required, RBC lysis buffer (BioLegend, cat#420301) was added for 5 minutes on ice before washing the buffy coat. The enriched mononuclear cells were washed twice with PBS containing 2% EDTA and centrifuged at 500 × g for 5 minutes. Cell count and viability were assessed using trypan blue hemocytometer counting.

For the addition of the transcription and translation cocktail, inhibitor stock solutions were reconstituted and stored as follows: Actinomycin D (Sigma-Aldrich, cat# A1410) was reconstituted in dimethylsulfoxide (DMSO, Thermo Fisher Scientific, cat# J66650.AE) at a stock concentration of 5 mg/ml and stored at -20°C, protected from light. Triptolide (Sigma-Aldrich, cat# T3652) was reconstituted in dimethylsulfoxide at a stock concentration of 10 mM and stored at -20°C, protected from light. Anisomycin (Sigma-Aldrich, cat# A9789) was reconstituted in dimethylsulfoxide at a stock concentration of 10 mg/ml and stored at 4°C, protected from light. All inhibitors were used within 14 days after reconstitution. The final concentrations used were 5 μg/ml of Actinomycin D (1:1000 from stock), 10 μM of Triptolide (1:1000 from stock), and 27.1 μg/ml of Anisomycin (1:368.5 from stock). The inhibitors were added to the whole blood within 5 minutes of the blood draw.

In the room temperature conditions, all reagents and steps of PBMC isolation, including centrifugation were performed at room temperature.

In the cold condition, the blood was placed on ice 5 minutes after the draw, all reagents and cells were kept on ice, with exception of the Ficoll that was room temperature and all centrifugations were performed at 4°C throughout the protocol.

### PBMC cultures with LPS treatment

PBMCs were plated at a concentration of 1 million cells/ml and treated with Lipopolysaccharides from *Escherichia coli* O111:B4 (Sigma-Aldrich, cat# L4391) diluted in RPMI media (Sigma-Aldrich, cat# R0883) with 10% FBS (Thermo Fisher, cat# 122484028) at 100ng/ml for 6 hours. Cells were either immediately exposed to LPS upon plating and harvested at the end of treatment or 18 hours after plating and harvested at the end of treatment.

### Time Post Thaw Experiments

For time post thaw experiments, cryopreserved PBMC cell vials were thawed briefly in a 37°C water bath, rinsed with pure FBS (Thermo Fisher, cat# 122484028), and quantified for plating at a concentration of 1 million cells in 1ml of RPMI media with 10% FBS (as above). After the specified time *in vitro*, cells were harvested, and RNA was isolated using the Qiagen RNeasy mini kit (Qiagen).

### RNA extraction

RNA was purified with the RNeasy mini kit (Qiagen) and quantified using a BioPhotometer plus model 6132 (Eppendorf).

### Real-time quantitative PCR

cDNA was prepared using superscript IV VILO (Thermo Fisher Scientific). Real-time quantitative PCR was performed using Taqman master mix (Applied Biosystems) on the QuantStudio7 (Applied Biosystems). Taqman primers (Applied Biosystems): *18*S (Hs99999901_s1), *ACTB* (Hs01060665_g1), *IL1b* (Hs01555410_m1), *IL6* (Hs00174131_m1), *DUSP1* (Hs00610256_g1), *JUNB* (Hs00357891_s1), *FOS* (Hs00170630_m1), *CCL4* (Hs99999148_m1), *CXCR4* (Hs00607978_s1). Delta Ct was calculated by normalizing the gene expression of each target mRNA to the expression levels of the housekeeping gene 18S that was found to be stable across experimental conditions.

### RNA sequencing

The NEBNext Ultra II directional RNA library prep kit for Illumina (New Englands Biolabs Inc., Ipswich, MA, USA) was used to prepare mRNA sequencing libraries, according to manufacturer’s instructions. Briefly, 150 ng of total RNA were purified using the NEBNext poly(A) mRNA Magnetic Isolation module (New Englands Biolabs Inc., Ipswich, MA, USA) and used as a template for cDNA synthesis by reverse transcriptase with random primers. The specificity of the strand was obtained by replacing the dTTP with the dUTP. This cDNA was subsequently converted to double-stranded DNA and end-repaired. Ligation of adaptors was followed by a purification step with AxyPrep Mag PCR Clean-up kit (Axygen, Big Flats, NY, USA), by an excision of the strands containing the dUTPs and finally, by a PCR enrichment step of 12 cycles to incorporate specific indexed adapters for the multiplexing. The quality of final amplified libraries was examined with a DNA screentape D1000 on a TapeStation 2200 and the quantification was done on the QuBit 3.0 fluorometer (ThermoFisher Scientific, Canada). Subsequently, mRNA-seq libraries with unique dual index were pooled together in equimolar ratio and sequenced for paired-end 100 bp sequencing on an Illumina NovaSeq 6000 at the Next-Generation Sequencing Platform, Genomics Center, CHU de Québec-Université Laval Research Center, Québec City, Canada. The mean coverage/sample was 12M paired-end reads.

### Bioinformatics

The raw paired-end fastq files obtained from RNA-sequencing were aligned to the human hg19 reference genome using STAR aligner (v2.6.1b)^26^. Next, the gene expression was obtained with FeatureCounts (v2.0.1)^27^. Group data was analyzed pairwise and normalized using the DESeq2 (v1.40.1)^28^ generalized linear model in R (v4.2.3). Differentially expressed genes (DEGs) were determined using cutoffs for p-value < 0.01 and an absolute log2FoldChange value of 0.58 (1.5 fold). Principal component analysis (PCA) and bar plots were made using ggplot2 (v3.4.2)^29^ in R. EnhancedVolcano (v1.18.0)^30^ was used to produce volcano plots. Pheatmap (v1.0.12)^31^ was used to produce heatmaps.

KEGG pathway and GO_Cellular component enrichment analysis on DEGs was performed using ShinyGO v0.77^32^, significance was set to FDR ≤ 0.05 and limited to the top 20 most significant categories for all analyses. For the hierarchical clustering tree summarizing the correlation among significant pathways, the pathways with many shared genes are clustered together. Bigger dots indicate more genes.

### Statistical analysis

Statistical analyses were performed using GraphPad Prism, Version 8 (GraphPad Software, Inc). Two-way ANOVA was used when more than two groups were compared with Sidak’s multiple comparison correction when appropriate. Error bars represent mean ± standard error of the mean (SEM). Significance level was set at p ≤ 0.05.

## Acknowledgments

We would like to thank the Next-Generation Sequencing Platform at the Genomics Center of the CHU de Québec-Université Laval Research Center for RNA sequencing and the CRCHUM UTMAB and C-BIG biobanks. We would also like to thank Christel Dias for coordinating blood draws. This study is funded by *Fondation Courtois and American Parkinson’s Disease Association (APDA) (969466)* as well as by the joint efforts of *The Michael J. Fox Foundation for Parkinson’s Research* (MJFF) and the *Aligning Science Across Parkinson’s* (ASAP) initiative. MJFF administers the grant ASAP 000525 on behalf of ASAP and itself. L.A. received scholarships from the Schlumberger Foundation and the Faculty of Medicine, University of Montreal. A.M. received a scholarship from FRQS/Parkinson’s Quebec. J.S and M.T are supported by a salary award from the Fond de recherche du Québec - Santé (FRQS).

## Author Contributions

Conceptualization, L.A., A.M., L.K.H., J.S., M.T; Methodology, L.A., A.M., L.K.H.; Investigation, L.A., A.M., L.K.H.; Formal analysis, L.A, L.K.H, M.L.; Writing – Original Draft, L.K.H., A.M.; Writing – Review & Editing, L.A, A.M. L.K.H., J.S., M.T.; Funding Acquisition, J.S, M.T.; Resources, M.-N.B., J.K., J.S., M.T.; Supervision, J.S., M.T.

## Data and code availability

Processed datasets created from this study are available in TableS1 and TableS2. Any additional information is available from the corresponding authors upon request.

## Declaration of generative AI and AI-assisted technologies in the writing process

During the preparation of this work the author(s) used OpenAI / ChatGPT in order to improve readability. After using this tool/service, the author(s) reviewed and edited the content as needed and take(s) full responsibility for the content of the publication.

## Supplemental Information

**Supplementary Figure S1: Adding TTi cocktail lyses red blood cells requiring addition of RBC lysis buffer for cell concentration determination**

**A** Schematic of the experimental plan for PBMC isolation with and without (+/-) RBC lysis buffer at RT (n=2/condition).

**B** Volcano plots of transcriptomic differences between PBMCs isolated at RT +/- RBC lysis buffer all without TTis showing 28 DEGs, with 25 down-regulated and 3 up-regulated.

**C** Volcano plots of transcriptomic differences between PBMCs isolated at RT +/- RBC lysis buffer all with addition of TTis showing 274 DEGs, with 149 down-regulated and 125 up-regulated.

**D** Schematic of experimental plan for PBMC isolation (+/-) RBC lysis buffer and cold.

**E** Volcano plots of transcriptomic differences between PBMCs isolated cold +/- RBC lysis buffer all without TTis showing 45 DEGs, with 35 down-regulated and 10 up-regulated.

**F** Volcano plots of transcriptomic differences between PBMCs isolated cold +/- RBC lysis buffer all with addition of TTis showing 21 DEGs, with 15 down-regulated and 6 up-regulated.

Error bars represent mean +/- SEM. Significance was set at p-value < 0.01 and an absolute log2FoldChange value of 0.58 (1.5 fold). Refer to TableS1 for raw transcriptomic data and TableS2 for complete DEG lists.

**Supplementary Figure S2: KEGG and GO analysis of down-regulated DEGs by cold processing or TTis show convergence on AP-1 complex.**

**A** KEGG pathway analysis of the 92 DEGs specifically down-regulated by cold processing predominantly identified ribosomal-related pathways.

**B** Gene ontology (GO) cellular compartment (CC) analysis of the 92 DEGs specifically down-regulated by cold processing revealed significant enrichment ribosomal-related processes.

**C** KEGG pathway analysis of the 216 DEGs down-regulated by addition of TTis predominantly identified cell contact and cell cycle-related pathways.

**D** Gene ontology (GO) cellular compartment (CC) analysis of DEGs down-regulated by addition of TTis revealed significant enrichment platelet and cytoskeletal-related processes.

Error bars represent mean +/- SEM. Significance was set at p-value < 0.01 and an absolute log2FoldChange value of 0.58 (1.5 fold). Refer to TableS1 for raw transcriptomic data and TableS2 for complete DEG lists.

**Supplementary Figure S3: KEGG pathway and GO enrichment analysis of DEGs between t0, t2, t24 hours post thaw.**

**A** KEGG pathway analysis of the 2256 DEGs down-regulated between t2pt and t0pt.

**B** Gene ontology (GO) cellular compartment (CC) analysis of the 2256 DEGs down-regulated between t2pt and t0pt.

**C** KEGG pathway analysis of the 2722 DEGs up-regulated between t2pt and t0pt.

**D** Gene ontology (GO) cellular compartment (CC) analysis of the 2722 DEGs up-regulated between t2pt and t0pt.

**E** KEGG pathway analysis of the 1041 DEGs up-regulated between t24pt and t0pt.

**F** Gene ontology (GO) cellular compartment (CC) analysis of the 1041 DEGs up-regulated between t24pt and t0pt.

**G** KEGG pathway analysis of the 1228 DEGs down-regulated between t24pt and t0pt.

Error bars represent mean +/- SEM. Significance was set at p-value < 0.01 and an absolute log2FoldChange value of 0.58 (1.5 fold). Refer to TableS1 for raw transcriptomic data and TableS2 for complete DEG lists.

