## Supplementary figures and images for "Comparative Analysis of Methods to Reduce Activation Signature Gene Expression in PBMCs"

### Figure S1_june.pdf

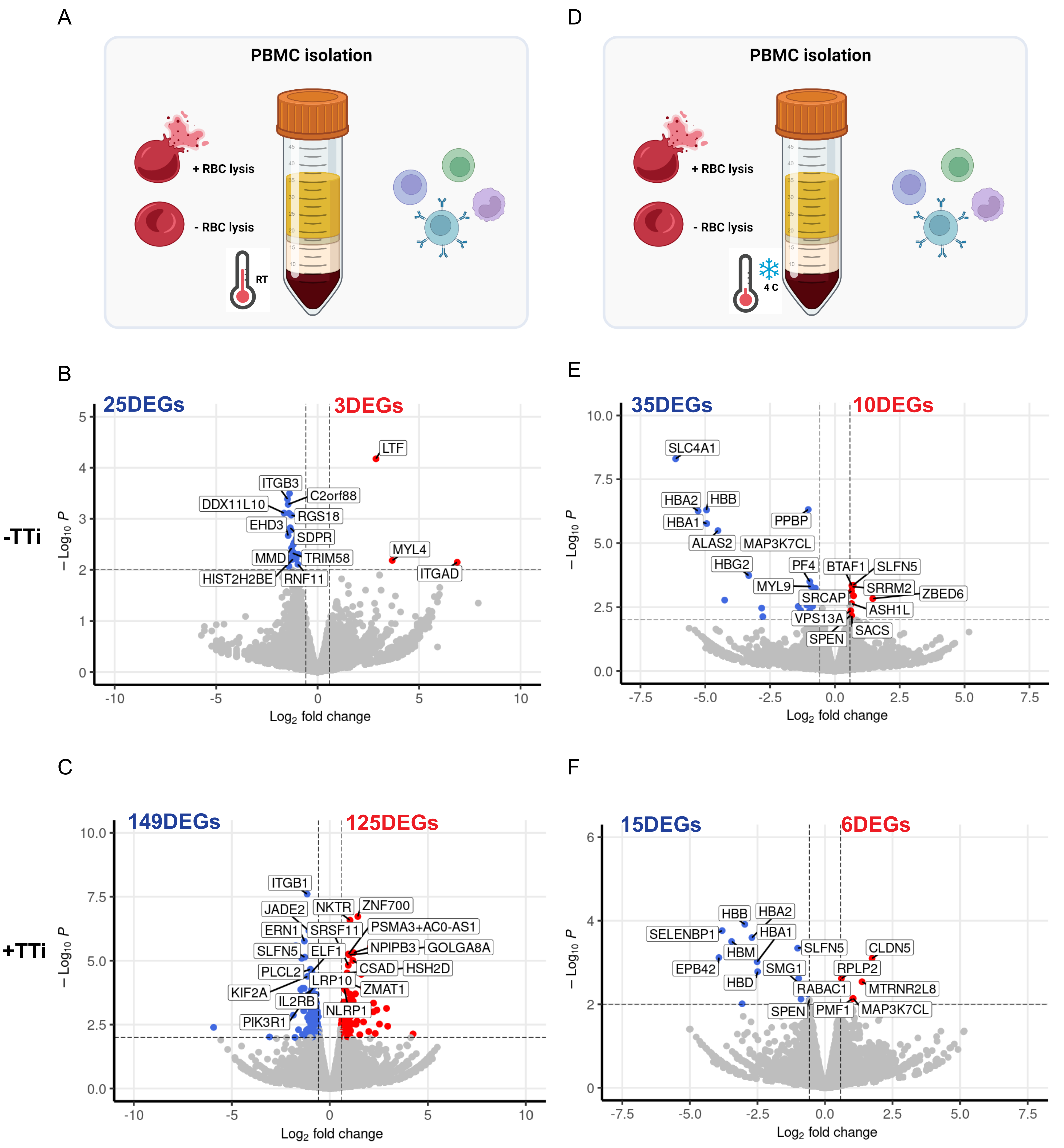

### Figure S2_june.pdf

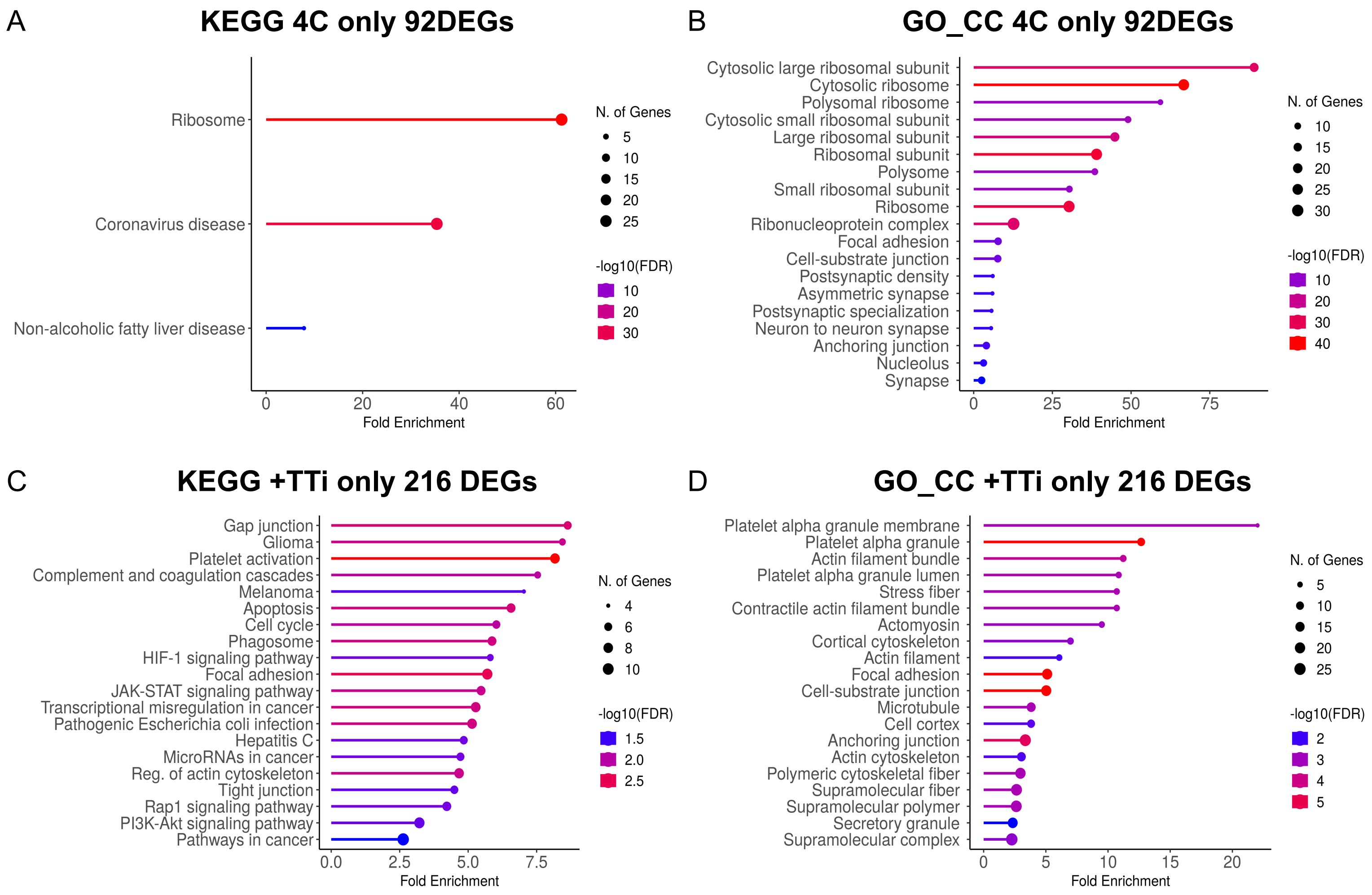

### FIgure S3-june.pdf

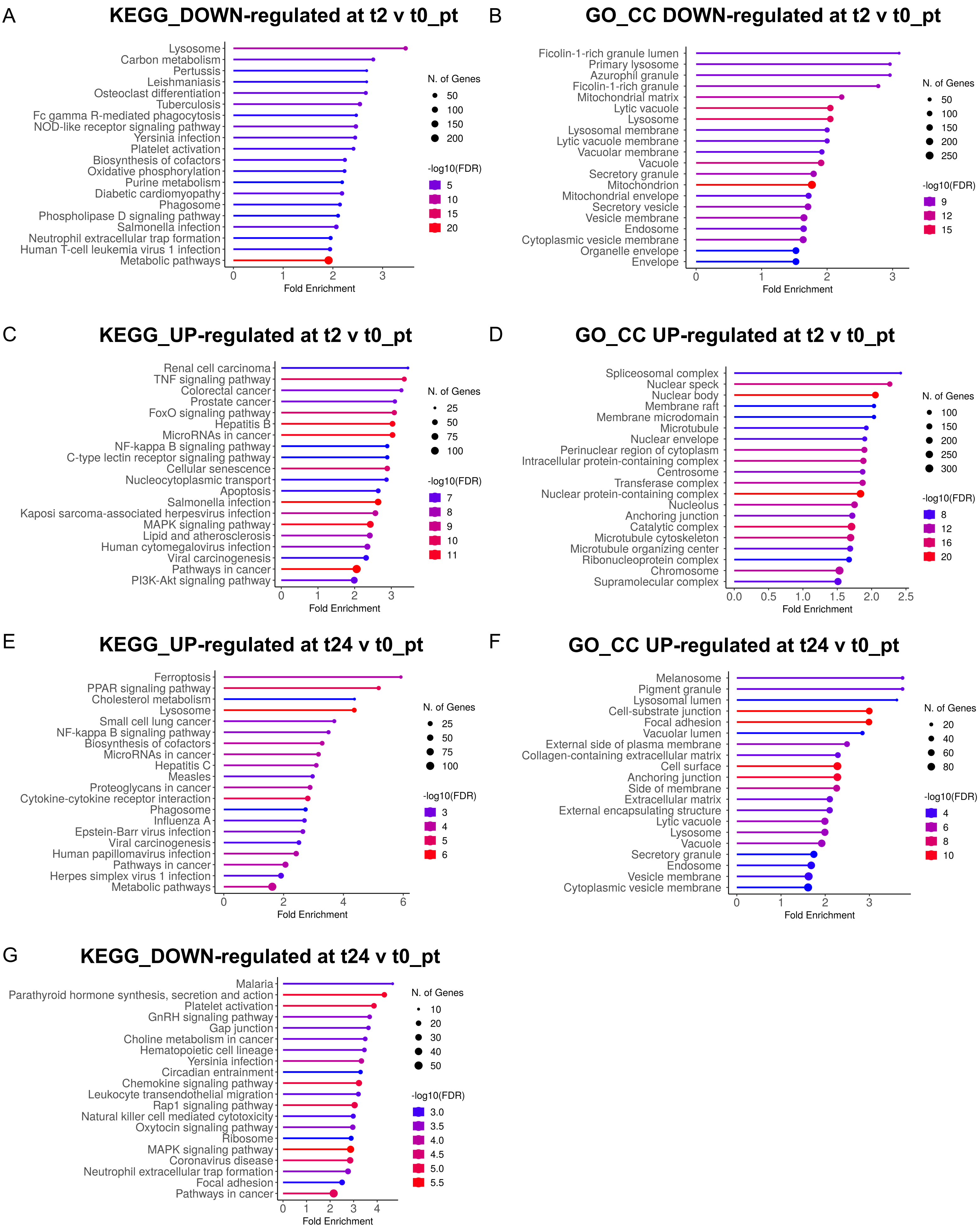
